# Endosymbiont diversity and evolution across weevil tree of life

**DOI:** 10.1101/171181

**Authors:** Guanyang Zhang, Patrick Browne, Geng Zhen, Andrew Johnston, Hinsby Cadillo-Quiroz, Nico Franz

## Abstract

As early as the time of Paul Buchner, a pioneer of endosymbionts research, it was shown that weevils host diverse bacterial endosymbionts, probably only second to the hemipteran insects. To date, there is no taxonomically broad survey of endosymbionts in weevils, which preclude any systematic understanding of the diversity and evolution of endosymbionts in this large group of insects, which comprise nearly 7% of described diversity of all insects. We gathered the largest known taxonomic sample of weevils representing four families and 17 subfamilies to perform a study of weevil endosymbionts. We found that the diversity of endosymbionts is exceedingly high, with as many as 44 distinct kinds of endosymbionts detected. We recovered an ancient origin of association of *Nardonella* with weevils, dating back to 124 MYA. We found repeated losses of this endosymbionts, but also cophylogeny with weevils. We also investigated patterns of coexistence and coexclusion.

## INTRODUCTION

Weevils (Insecta: Coleoptera: Curculionoidea) host diverse bacterial endosymbionts. In his monumental book, “Endosymbiosis of Animals with Plant Microorganisms”, Buchner (1965) (p. 160) remarked that “nowhere else aside from the cicadas [Hemiptera: Auchenorrhyncha and Sternorrhyncha] do so many symbiotic sites exist as in this insect family [Curculionidae sensu lato]”. (citation not found: Nardon&Grenier) (p. 181) listed nine morphological types of endocytobiosis (a.k.a, bacterial endosymbiosis) in Curculionidae (excluding Scolytinae, therein as Scolytidae). They lamented on the ubiquity of bacterial endosymbionts in weevils and suggested that “most if not all Curculionids harbor endocytobiotic bacteria” (p. 189). Despite this long-standing realization, the diversity, evolution and biology of bacterial symbionts of weevils have been poorly documented or understood, especially if compared to Auchenorrhyncha and Sternorrhyncha (“Homoptera”). With 62,000 described species and 220,000 estimated to exist (Oberprieler 2007), weevils are a mega-diverse group of insects and represent a premier system for studying bacterial symbionts.

The astounding diversity of weevils strongly contrasts with the rather small number of species that have been studied for bacterial symbionts. Early research using morphological techniques examined only some 30 genera of weevils (Buchner 1965), and these typically did not reveal the taxonomic identities of the symbionts. The first molecularly characterized endosymbiont from weevils was SOPE, or “Sitophilus oryzae Primary Endosymsbiont”. This was later named as “*Candidatus* Sodalis pierantonius” (the qualifier *Candidatus*, indicating that the species name is provisional [Murray 1995], is only used at the first occurrence of a name and will be omitted from subsequent occurrences). *Sodalis* was previously placed in Enterobacteriaceae, which has been split into several families (Adeolu 2016), resulting in its new placement in Pectobacteriaceae of the order Enterobacteriales, in the class Gammaproteobacteria. Sodalis was first described from tsetse flies (Glossinidae, Diptera) (Dale 1999) and has also been found infecting various insects including Aphididae, Cercopidae, Cicadellidae, Pentatomidae, Scutelleridae, (mostly plant sucking insects; Hemiptera), Apidae, Megachilidae (bees; Hymenoptera), Philopteridae (lice; Phthiraptera), and Cerambycidae (longhorn beetles; Coleoptera). Lefevre (2004) described “*Candidatus* Nardonella” (Enterobacteriaceae) from a group of dryophthorine weevils (Dryophthorinae/-idae). Subsequent studies found *Nardonella* from the weevil subfamilies Brachycerinae/-idae (Erirhininae/-idae), Cyclominae, Entiminae and Molytinae (Hosokawa 2010, Hosokawa 2015, Huang 2016, Hirsch 2012, Merville 2013, Conord 2008, White 2015). To date Nardonella has not been found from other insects and its association with weevils is unique among insects. In a series of studies, Toju et al. (2009); Toju et al. (2013); Toju et al. (2010) discovered and documented “*Candidatus* Curculioniphilus” (Enterobacteriaceae) from several genera within the Curculioninae, namely, *Curculio*, *Shigizo*, *Archarius*, *Koreoculio* and *Carponinus*. *Curculioniphilus* is also a weevil-specific bacterial lineage. Besides *Sodalis*, *Nardonella* and *Curculioniphilus* other bacterial endosymbionts such as *Wolbachia*, *Rickettsia* (Alphproteobacteria), *Spiroplasma* (Mollicutes), *Klebsiella pneumonia* (Gammaproteobacteria) have been previously found from weevils. Additionally, several weevils of (*Trigonotarsus* and *Diocarandra*) are associated with several distinct and unnamed endosymbionts. Here we provide a succinct summary of the published findings.

Previous studies on weevil endosymbionts were limited in the taxonomic scope of weevils. The phylogenetic space or the tree of life of weevils thus remains poorly sampled. This study represents the first attempt to gather a large taxonomic sample of weevils that represents a comprehensive phylogenetic diversity, albeit sparsely, to investigate the diversity and evolution of their associated endosymbionts.

## MATERIAL AND METHODS

### Insect taxonomic sampling and specimen vouchering

We performed DNA extractions and PCR amplifications on a total of 244 beetle specimens, which represented five superfamilies (Chrysomeloidea: Cerambycidae, Chrysomelidae, Disteniidae; Cleroidea: Cleridae, Melyridae, Trogossitidae; Cucujoidea: Coccinellidae, Erotylidae; Curculionoidea: Anthribidae, Attelabidae, Brentidae, Curculionidae, Erirhinidae; Tenebrinoidea: Tenebrionidae). Within the Curculionoidea, four families – Anthribidae, Attelabidae, Brentidae, and Erirhinidae – were studied for the first time for their bacterial symbionts using a molecular method. Within the family Curculionidae, 17 subfamilies were sampled, 13 for the first time.

We assigned a morpho-species code starting with “BEP” (Bacterial Endosymbionts Project) to each species (e.g., BEP 123). For each morpho-species, the specimen used for DNA extraction, whether dissected or intact, was designated as the primary voucher, and additional conspecific specimens, if available, were designated as secondary vouchers. All voucher specimens were deposited at the Arizona State University Hasbrouck Insect Collection (ASUHIC) and databased using the SCAN (Symbiota Collections of Arthropods Network) database (Gries et al., 2014) and the specimen records are accessible online (http://symbiota4.acis.ufl.edu/scan/portal/index.php). Morpho-species code and voucher status were recorded in the ‘notes’ field (e.g., BEP 123, primary voucher). Of the 244 beetle specimens, 124 (51%) generated usable PCR products (see below), 115 of which were weevils and nine other beetles.

### DNA extraction and PCR amplification of bacterial 16s

For species with two or more conspecific specimens, we dissected the abdomen and isolated the gut contents of one specimen, which was used for DNA extraction. For singleton species with only one specimen available, we used the entire specimen for DNA extraction, without gut dissection. Of the specimens that generated usable PCR products, 103 were subjected to gut dissection and 21 were extracted from whole bodies. The dissected gut contents were subjected to one minute of (twice of 30 seconds) bead beating with bead beater before used for DNA extraction. DNA extraction was performed with the Qiagen Blood and Tissue DNeasy Kit.

We performed two rounds of PCR amplifications. We first used regular primers that were not barcode-tagged (515F: 5’-GTGYCAGCMGCCGCGGT-3’ and 909R: 5’-CCCCGYCAATTCMTTTRAGT-3’), to amplify a 400 bp (base pair) fragment of the gene 16s. Gel electrophoresis of the PCR products recovered bands consistently for the negative control, probably due to reagent contamination, although usually weaker than the bands of the actual samples. Samples that had bands stronger than the negative controls were then used for the second round of PCR amplification. The primers used in the second round of PCR amplification were barcode-tagged at the 5’ end. Each barcode consists of six nucleotides and differs from one another by two nucleotide positions. Altogether 138 samples went through two rounds of PCR, of which 124 were successful in both rounds. The PCR products of those samples were subsequently sequenced and are here called usable PCR products.

### Post-PCR processing and sequencing - gel extraction, quantification, normalization, library preparation and Illumina sequencing

Two PCR bands were obtained for most samples. This was unexpected and led us to investigate the specificity of the primers. It was determined that the primers align near perfectly with insect 18s sequences and define a region longer than the target bacterial 16s fragment. Therefore, we deemed that the smaller fragment represented the bacterial 16s gene. Gel extraction was performed with the Promega Wizard^®^ SV Gel and PCR Clean-Up System to extract and purify bacterial 16s bands. The purified PCR products were Quantified with a NanoDrop spectrophotometer. Normalization was performed using the SequalPrep™ Normalization Plate Kit by Thermo Fisher Scientific. This created a pooled sample containing 124 PCR products. This was further subjected to Illumina library preparation performed by the “DNASU Next Generation Sequencing Laboratory” (http://dnasusequencing.org/). Paired-end sequencing was performed on a single Illumina MiSeq lane.

### Read processing – demultiplexing, quality filtering, and chimera removal

A python script was used to demultiplex paired reads, trim barcode and primer sequences and to perform quality control. The script required exact matches to both barcodes and for each of the forward and reverse primer sequences to be present with a maximum of three mismatches to the degenerate primer sequences in order for paired reads to be binned into a sample. The quality trimming removed the barcode and primer sequences and trimmed the 3’ end of the reads from the first point where a Phred score of less than 2 was encountered. If, after trimming, either of a pair of reads was shorter than 200 nt, the read pair was discarded.

After demultiplexing, read pairs were assembled using the program PEAR (Zhang et al., 2014). Following read assembly, reads were quality filtered using the ‘maxee’ parameter of usearch (Edgar, 2010) to remove any reads with more than one expected error. We obtained a dataset of 11,396,976 sequences. The assembled reads were dereplicated and singletons were removed from the dataset using VSEARCH (Rognes et al., 2016) (https://github.com/torognes/vsearch). Reference based and de novo chimera detection was performed using vsearch’s “uchime_ref” and “uchime_denovo” functions respectively on sequences within each sample. It would be computationally prohibitive to perform chimera detection on the pooled dataset. The abskew parameter was set to 1.5 for the de novo chimera detection and a customized version of the May 2013 version of the greengenes 16s database, modified to include known endosymbiont lineages, was used for reference based chimera detection (see “supplementary method description” in Supporting Information). Sequences flagged as chimeric by either method, de novo or reference based, were discarded. The quality-filtered, dereplicated, chimera-removed dataset contained 282,587 sequence reads.

### Phylogenetic identification of endosymbiont sequences

We devised a method termed “phylogenetic identification of next generation sequences” (PINGS) to perform taxonomic assignments of 16s amplicon sequences. We resorted to this method because an initial analysis in QIIME using a distance-based taxonomic assignment method failed to place 45% of our sequence reads at the genus level or below. Moreover, weevil-specific endosymbionts such as Nardonella and Curculioniphilus were not recovered in that analysis. The PINGS method consists of five steps and a succinct description is provided here: sequencing filtering, scaffold phylogeny reconstruction, symbiont screening, phylogenetic verification, and taxonomic assignment. Sequence filtering entailed the use of USEARCH to filter the dereplicated, chimera-removed dataset to retain only sequences with 100 and more reads. This resulted in a dataset of 4,217 sequences and we call this the “100 plus” dataset. The next step, scaffold phylogeny reconstruction, we started with generating a genus-level reference 16s dataset by filtering the “custom greengenes 16s reference dataset” to retain only sequences with genus names (many sequences in the greengenes dataset do not have names at the genus level). This dataset (1,209 sequences) was combined with the 100 plus dataset and together they were aligned using MAFFT version 7 (Katoh 2009, Katoh 2013) to generate an alignment of 5426 sequences. A maximum likelihood phylogeny was reconstructed using RAxML (Stamatakis 2014) scaffold phylogeny. During the next step, symbiont screening, we inspected the scaffold phylogeny to screen for “provisional endosymbionts”, using Dendroscope 3 (version 3.5.7) (Huson 2012). A sequence was considered as a “provisional endosymbiont” if it either clustered with reference endosymbionts or appeared to be highly divergent from its closest relatives and also was highly abundant (read number greater than 1000). During initial data exploration, we observed that different samples shared identical sequences, probably a result of mismatching barcodes (Sinclair et al., 2014), and also that the true sequence had the highest read number, often 10 times or greater than those of the false positives. We excluded these false positive sequences. Because the taxonomic sampling of symbionts and their closely related taxa in the reference 16s dataset was not comprehensive, the identities of the provisional endosymbiont sequences would need to be further tested, which constitutes the final step called phylogenetic verification. Here, the provisional endosymbiont sequences were queried against Genbank using BLAST, and the sequences of top matches representing different species or genera were downloaded. These included both determined sequences (to genus or species) and undetermined sequences that were proposed to be insect symbionts in the study that report them. Besides, a taxonomically broad sample of related sequences (both symbionts and non-symbionts) were downloaded from Genbank. Finally, these sequences (BLAST matches and separate downloads) were aligned with the provisional endosymbiont sequences and maximum likelihood phylogenetic analyses was performed. In the last step, taxonomic assignment, we inspected each phylogeny and assigned taxonomy to the endosymbiont sequences. For each sequence we made a short statement regarding its phylogenetic relationship to reference bacteria, such as “part of Wolbachia clade” or “sister to Prevotellaceae”. Assignment to genus or species was achieved for most sequences, but not possible for some sequences and this is because they formed a sister relationship to a group of genera or appeared to be highly divergent from their closest relatives. Those were assigned to the next higher taxonomic rank (family or order).

### Tests for correlation or coexistence

A Spearman’s rank correlation test was performed to test for direction and strength of correlation between pairs of major endosymbiont groups and the results were visualized using the R packages corrplot (Wei 2016) and Hmisc(Jr 2017). Chi-square tests, implemented in R statistical environment, were used to examine if the occurrences of endosymbionts were non-randomly distributed in i.e., Nardonella-free versus Nardonella-positive samples, or if Nardonella had an impact on the occurrences of other symbionts.

### Terms and definitions

Reads are unprocessed Illumina sequences and are redundant. Abundance of a sequence is the number identical reads representing that sequence. A sample refers to the total collection of sequences derived from a single specimen. We use the term “kind of endosymbiont” as the basic taxonomic unit here. Most of the time this corresponds to a genus, e.g., *Spiroplasma*, but sometimes a group of inadequately determined endosymbiont sequences (e.g., Porphyromonadaceae symbiont of weevil; Table 1). An occurrence refers to the observation of one kind of endosymbiont in a sample. Co-existence is defined as the observation of two occurrences of endosymbionts in the same sample.

**Table 1.**
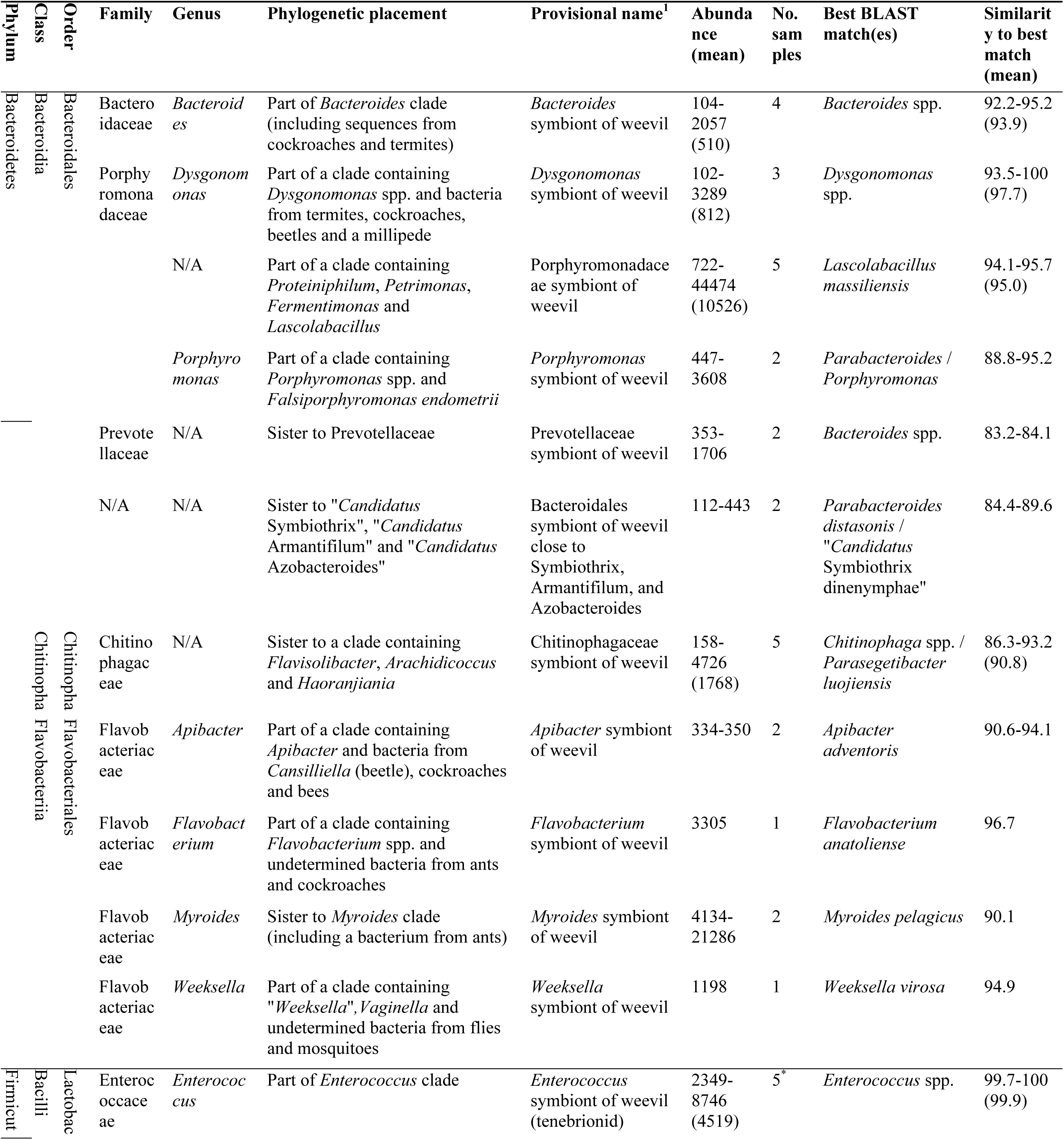
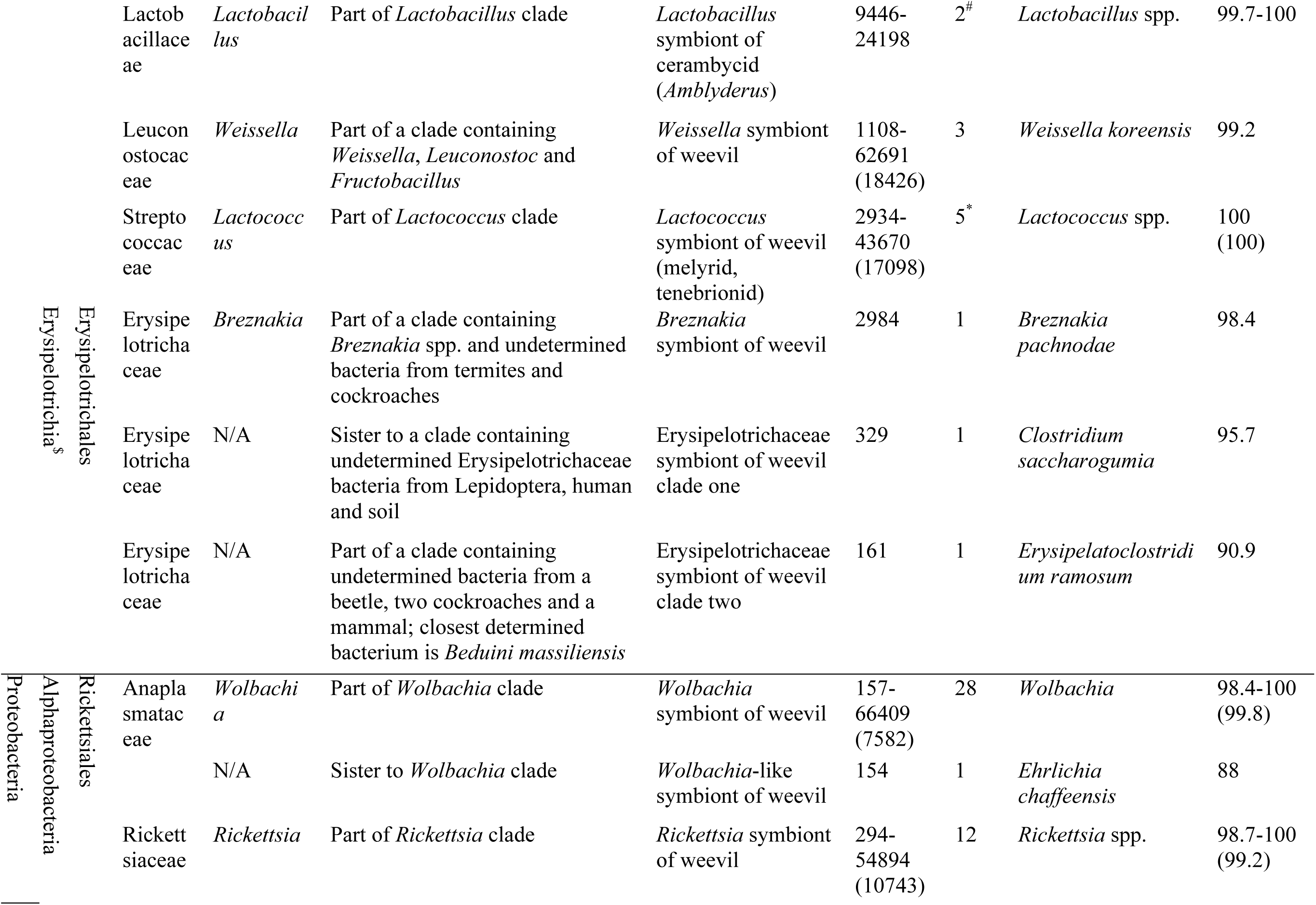
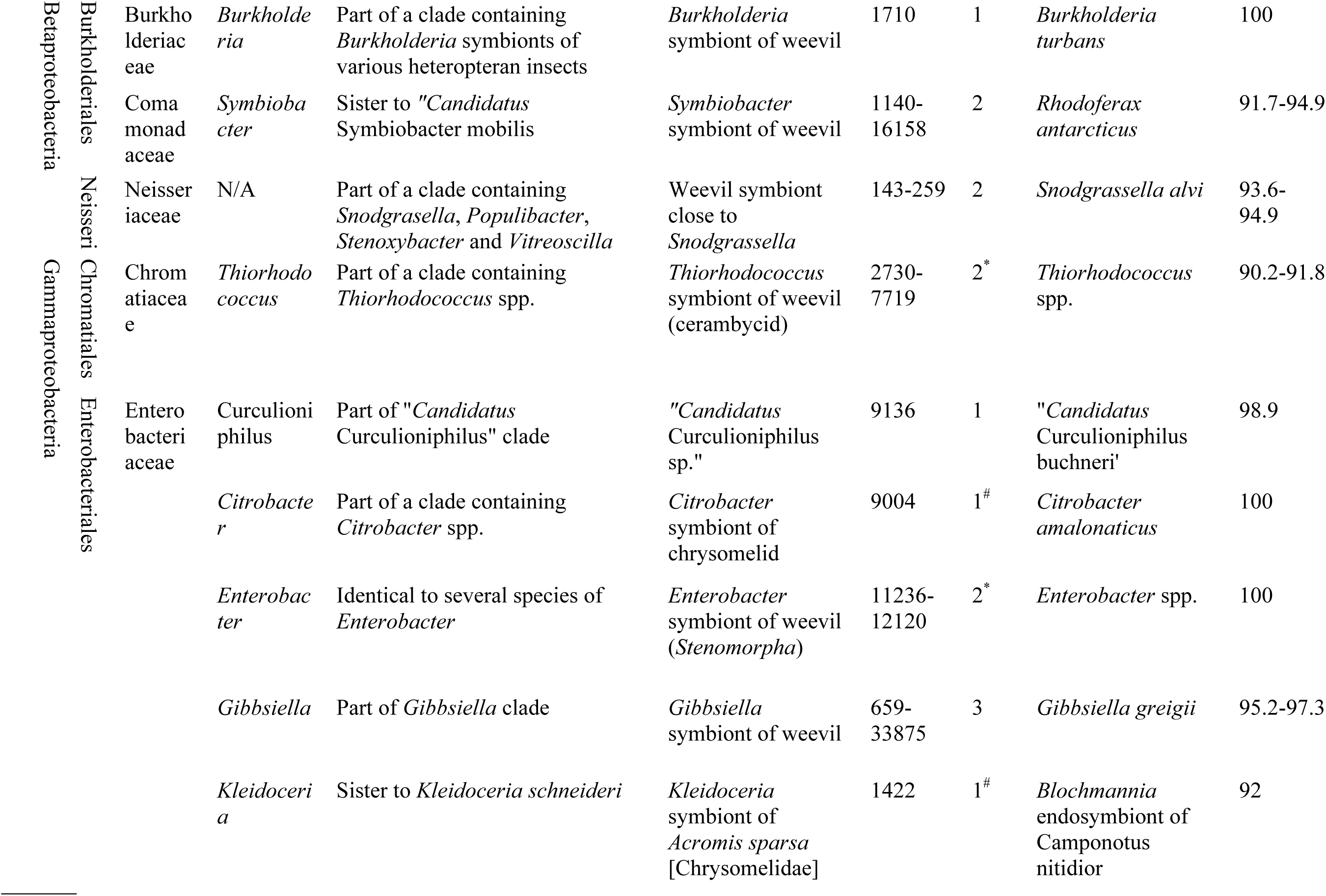
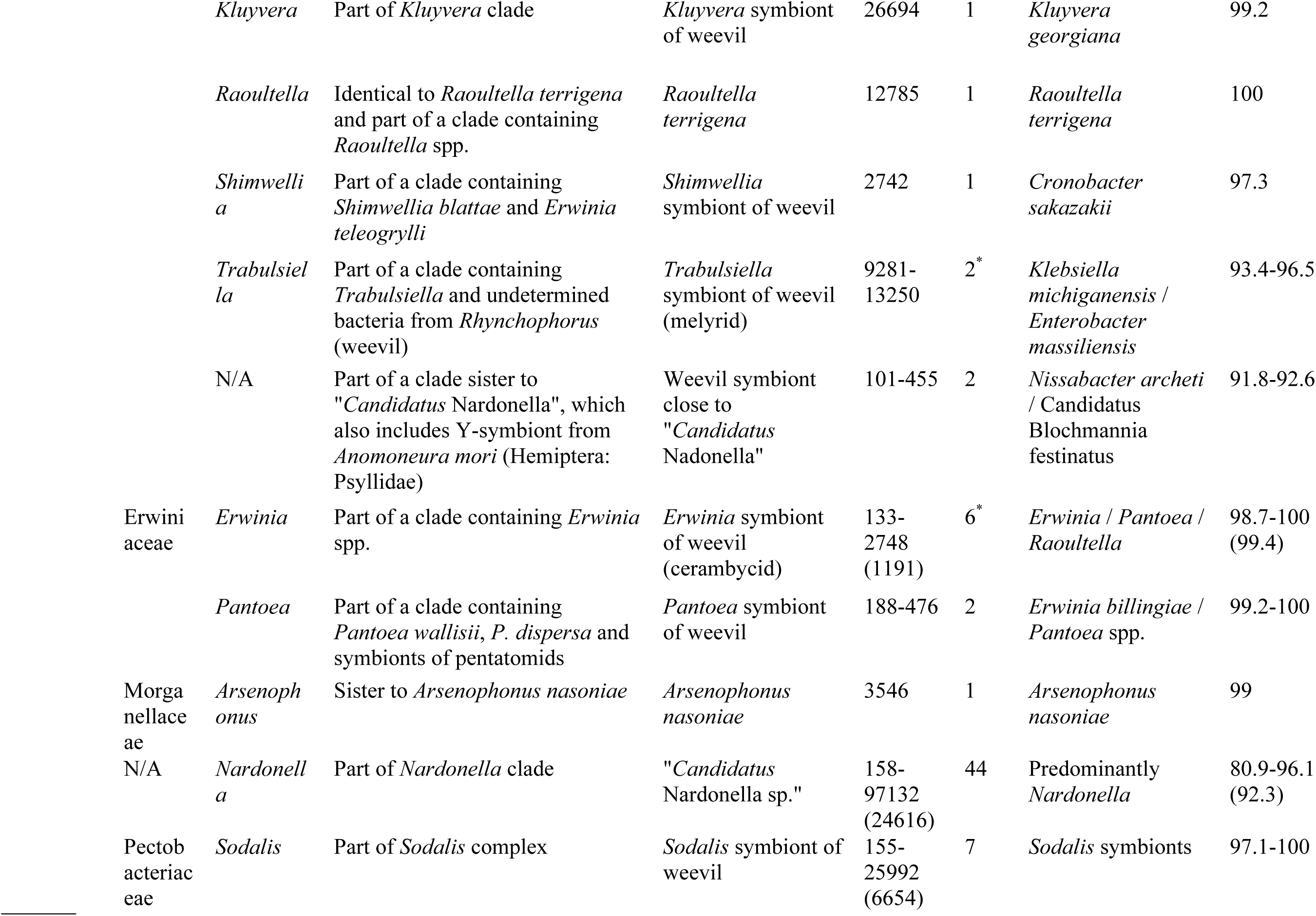
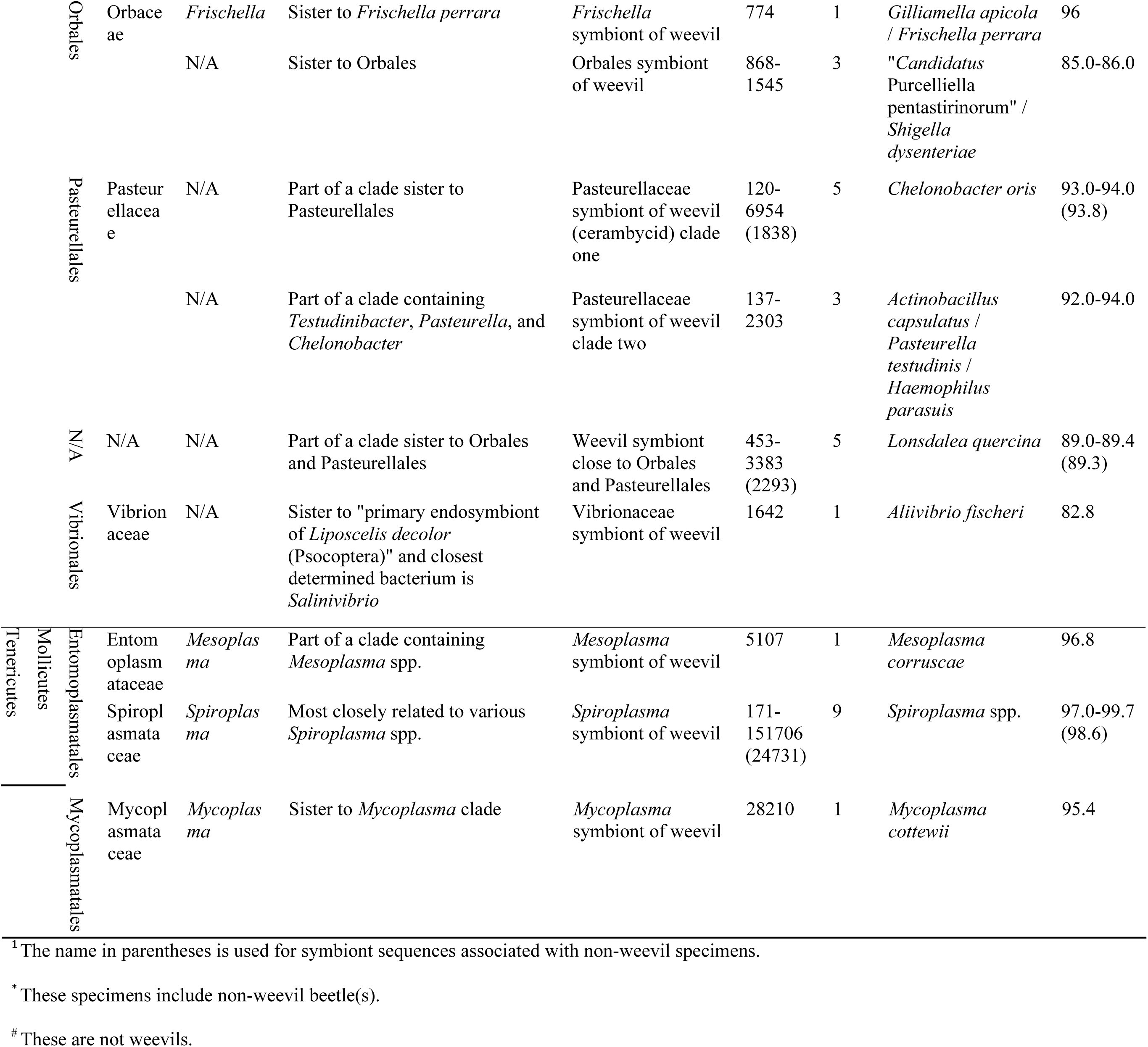
Taxonomic assignments, abundance and BLAST matches of symbiont sequences of weevils (Curculionoidea) and other beetles. Higher-level classification primarily follows that of NCBI Taxonomy. Phylogenetic placement is a short statement that describes the phylogenetic relationship of undetermined sequences to reference bacteria on a given phylogeny (reconstructed using 16s sequences and maximum likelihood method in this study). An existing taxonomic name is used if it captures the phylogenetic relationship, but an informal name is created to accommodate a “novel” (previously unnamed or unknown) relationship. Nearly all “provisional names” carry a modifier “symbiont of weevil” because in the absence of a well-curated database of insect bacterial symbionts, this will facilitate data discovery and communication. Abundance refers to the read number of a sequence. Only determined sequences (to species or genus for certain symbionts) in Genbank are considered for best BLAST matches. Similarity is a percentage value. Mean values of abundance and similarity are provided only if the number of sequences is equal to four or greater. In ‘provisional name’ column, add a note to indicate if the bacteria is previously known as an insect symbiont.

## RESULTS & DISCUSSION

### Sequence data profile

The demultiplexed, unfiltered data set contained a total of 11,396,976 reads. The number of reads per sample ranged from 4,483 to 457,087 (mean = 91,911, median = 64,972, stdev = 86,430). The dereplicated, chimera-filtered data set contained 5,820,560 reads, which represented 282,587 unique sequences. The total number of reads per sample in this data set ranged from 845 to 289,425 (mean = 46,940, median = 29,290, stdev = 50,786; Fig. 1a).

**Figure 1.**
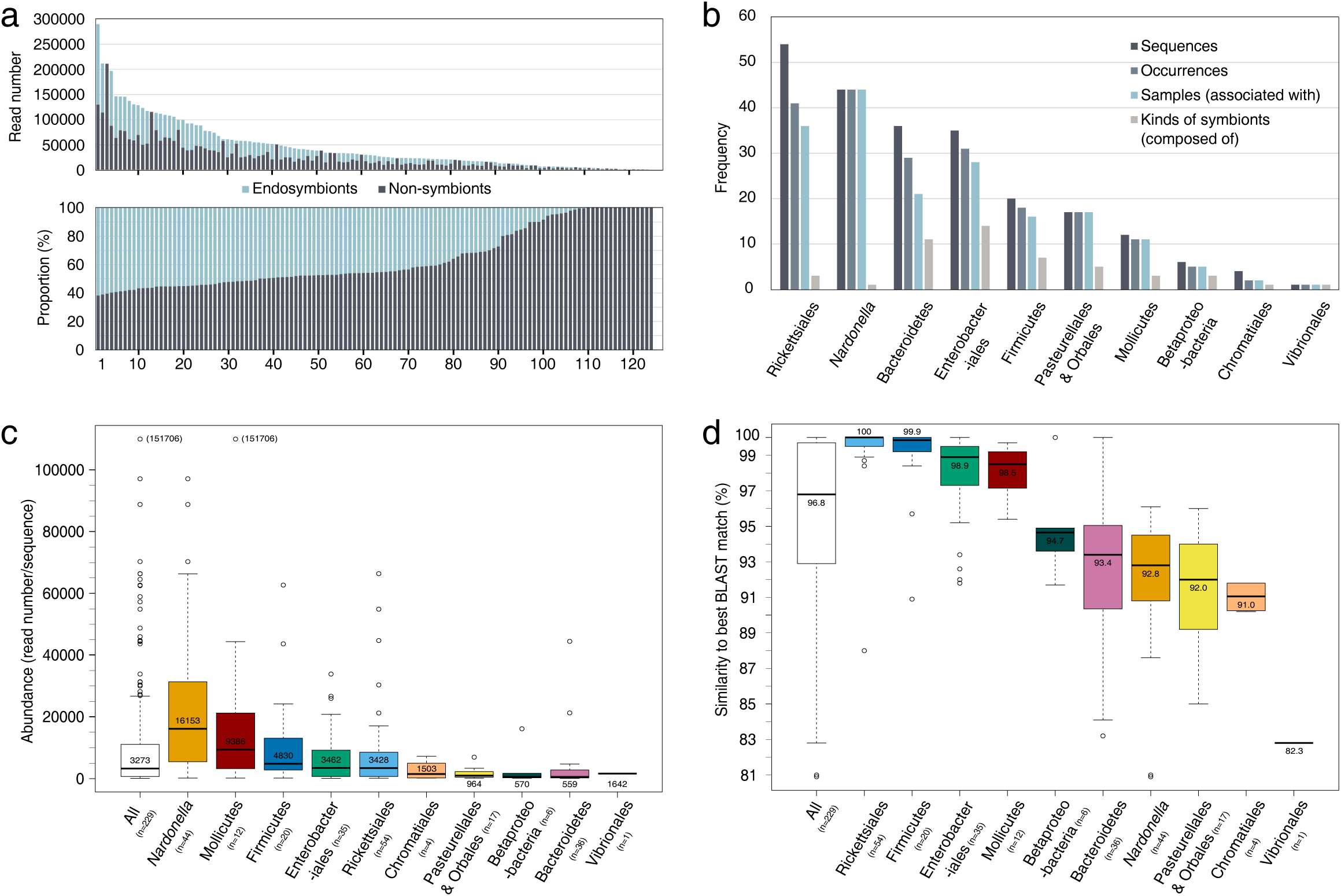
16s sequence data profile. (a) Total read numbers (upper) and proportions of endosymbiont and non-symbiont reads (lower) across 124 samples. Each bar represents a sample. The read numbers are based on the dereplicated, chimera-filtered data set. Bars in upper and lower charts do not match. (b) Numbers of sequences, occurrences, associated samples, and kinds in 10 major groups of endosymbionts. (c) Boxplot of abundance values of 10 major groups of endosymbiont sequences. The upper and lower borders of a box correspond to the first and third quartiles, and each box includes the middle 50% values in a group, or the interquartile range (IQR). The horizontal line inside a box is drawn at median, and its value indicated above the line. The upper or lower fence (not drawn) is a value that is 1.5xIQR above or below the third or first quartile. The horizontal lines at the ends of whiskers (i.e., vertical dotted lines) represent the maximum or minimum observation among all values in a group (when outliers are absent) or the observation that is closest to the upper or lower fence. Unfilled circles are outliers, which are values greater or smaller than the upper or lower fence. (d) Boxplot of similarities of endosymbiont sequences to best BLAST matches.

### Bacterial endosymbionts—phylogenetic identity, abundance and genetic divergence

We determined that 41.9% (2,436,371) of the dereplicated reads, representing 229 unique sequences, came from bacterial endosymbionts. Of the 124 samples sequenced, 109 (87.9%) were infected with at least one endosymbiont, 101 of which are weevils. For simplicity of presentation, we do not distinguish weevils and other insects when describing the results, unless it is otherwise stated. In a majority (61% or 76/124) of the samples, endosymbionts comprised greater than 40% of the reads in a sample (Fig. 1a). The non-symbiont reads also included those that were variants (generated from PCR or sequencing errors) of the most abundant endosymbiont sequences, so the actual proportion of endosymbionts in each sample would be somewhat higher than shown in Fig. 1a. All endosymbiont sequences comprised 199 occurrences (Fig. 1b). A total of 49 kinds of endosymbionts were identified based on the PINGS method, 46 of which were associated with weevils and 3 exclusively with non-weevil beetles (Table 1). These were placed in 27 families, 15 orders, 9 classes, and 4 phyla (Table 1). Composed of 28 kinds of endosymbionts, Proteobacteria showed the highest diversity among our sequences, followed by Bacteroidetes (11), Firmicutes (7) and Tenericutes (3). Thirty five (71%) kinds of endosymbionts were placed to a genus, but 14 (29%) could not be assigned to any genera and instead were placed at the family or order level (Table 1). These sequences diverged significantly from their closest relatives, and were 82.8-95.7% similar to their best BLAST matches. The most common five kinds of endosymbionts, in terms of the number of samples associated, are Nardonella (associated with 44 samples), Wolbachia (28), Rickettsia (12), Spiroplasma (9) and Sodalis (7).

We classified the 49 kinds of endosymbionts into 10 major groups as described in the following. Bacteroidetes, Tenericutes (represented by Mollicutes in this study) and Firmicutes constituted 3 groups as each is a different phylum. The remaining 7 groups all belong to the phylum Proteobacteria. First, these were separated by class, which resulted in recognizing Rickettsiales (the only order of the class Alphaproteobacteria recovered in this study) and Betaproteobacteria as 2 groups and Gammaproteobacteria containing the remaining 5 groups. Nardonella was considered as its own group as this is the most widely present endosymbiont and also one of the focal endosymbionts of our study. Enterobacteriales (treated as excluding Nardonella throughout the text), Pasteurellales & Orbales (P & O), Chromatiales and Vibrionales, four different order-level taxa, were treated as different groups. This grouping scheme allowed us to examine patterns of endosymbiont diversity, divergence and co-existence at a level that is reasonably granular yet cohesive. For each major group of endosymbionts, the number of occurrence is as follows: Nardonella (44), Rickettsiales (41), Enterobacteriales (31), Bacteroidetes (29), Firmicutes (18), P & O (17), Mollicutes (11), Betaproteobacteria (5,) Chromatiales (2), and Vibrionales (1).

The abundance of the endosymbiont sequences ranged from 101 to 151,706 (median = 3,273, mean = 10,639, stdev = 19,027; Fig. 1c). About 30% (69/229) of the sequences exhibited relatively low abundance (100-1000), 44% medium to high abundance (1000-10,000), and 26% very high abundance (>10,000). Nardonella showed the highest median abundance (16,153), followed by Mollicutes and Firmicutes, and the Bacteroidetes the lowest (Fig. 1c). All but two groups (P & O and Vibrionales) contained at least one sequence with abundance greater than 10,000. The highest abundance (151,706) of any sequences was found in a *Spiroplasma* (Mollicutes) endosymbiont (Table 1, Fig. 1c).

Half of the sequences (115/229) shared less than 97% similarity to their respective best BLAST matches, and those were pertinent to 30 kinds or 61% of the endosymbionts (Table 1, Fig. 1d). The median similarity across all endosymbionts was 96.8% (Fig. 1d). All the sequences of Nardonella, P & O, Chromatiales and Vibrionales showed less than 97% similarity to their best BLAST matches. The BLAST matches of 76% (174/229) sequences were congruent with the phylogenetic placements of the endosymbionts at the genus level (regardless of percentage similarity) (Table 1). The lowest similarity (80.9%) was found in a sequence of Nardonella. Interestingly, Only two kinds of endosymbionts, Dysgonomonas (Bacteroidetes) and Gibbsiella (Eneterobacteriales), contained sequences that showed similarity both above and below 97% to their best BLAST matches (Table 1).

### Nardonella—origin, repeated losses and co-evolution

We recovered the weevil-specific endosymbiont Nardonella from Brentidae, more specifically, the subfamily Apioninae, for the first time (Fig. 2). Brentidae is the sister group of Curculionidae and together they are here referred to as the “BC clade”. We thus provide evidence to support that the origin of Nardonella association with weevils dates back to 124.5 MYA (million years ago), or the Early Crestaceous. Nardonella was not detected from Attelabidae (n=5) or Anthribidae (n=3; 1 sample not shown in Fig. tree as no symbionts were recovered), the other two weevil families examined. Within the Curculionidae, Nardonella was newly recovered from Baridinae, Ceutorhynchinae and Conoderinae, and from many genera previously not sampled in Cryptorhynchinae, Dryophthorinae, Entiminae, and Molytinae. Nardonella was entirely absent from Lixinae, Curculioninae and Cossoninae as sampled in this study. While its distribution across the weevil tree of life is scattered from Brentidae to various clades of Curuclionidae, a dense concentration of Nardonella is seen in two clades, a Cryptorhynchinae clade and a core Baridinae-Molytinae clade. Across the phylogeny, the absence of Nardonella is sporadically distributed, rather than concentrated in a certain part of the tree. We consider this pattern, i.e., widespread presence of Nardonella and sporadic absence across the phylogeny, as evidence for recurrent losses of Nardonella-weevil association. An alternative explanation would be multiple independent acquisitions from other insects or the environment, but this is improbable as Nardonella is not known to occur in any insects or the environment. There would be no source for external acquisition. Also, the highly divergent 16s genes of Nardonella also lend support to the ancient origin, accompanied with recurrent losses explanation.

**Figure 2.**
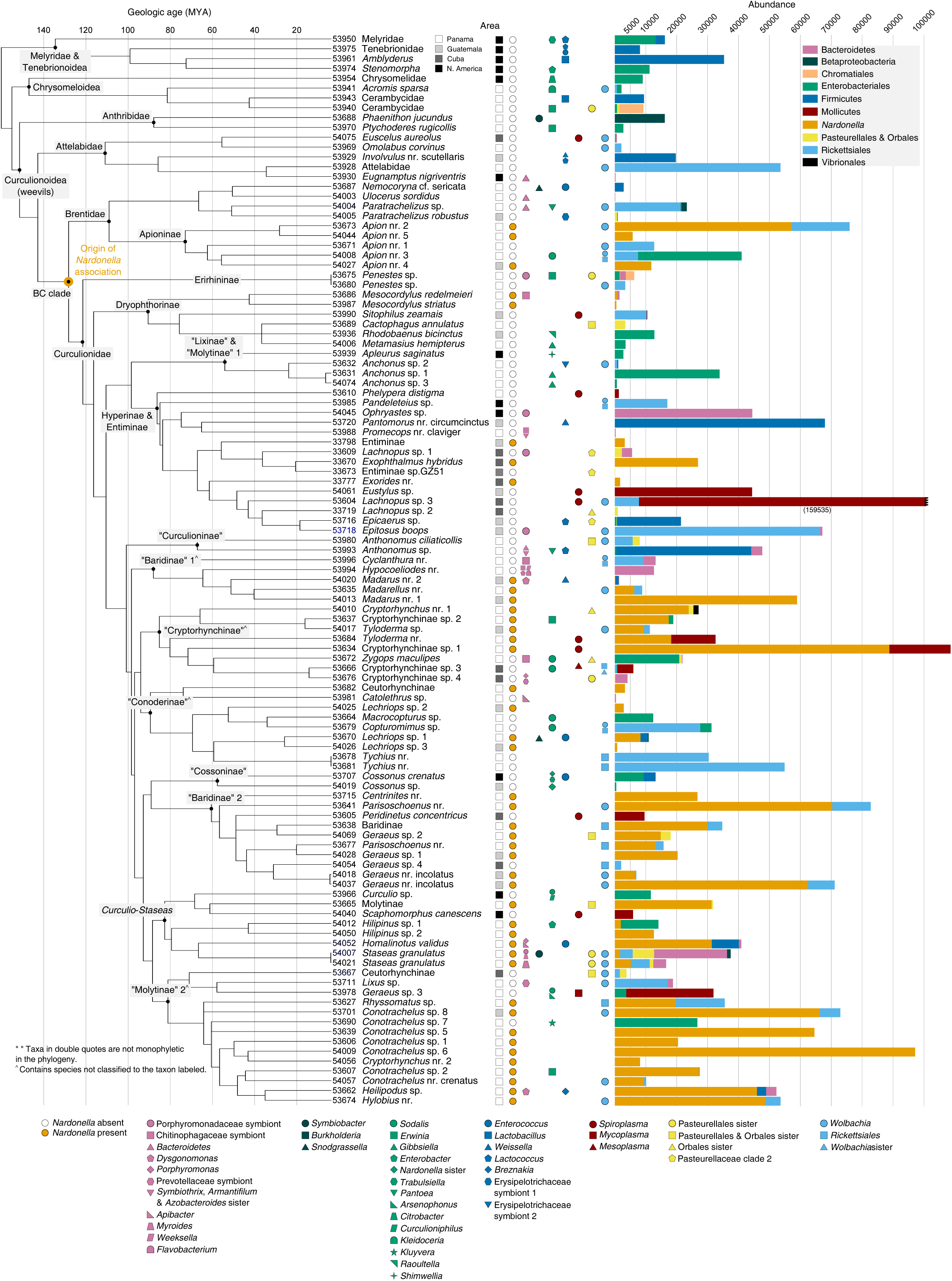
Endosymbiont diversity and sequence abundance are mapped onto a time-calibrated weevil phylogeny. The length of a bar is proportional to the total abundance an endosymbiont group (which would include one or several symbionts). Within a group, specific kinds of endosymbionts are denoted by different shapes. Taxa in double quotes are not monophyletic in the current study. ^^^Clade contains taxa previously not classified to the labeled taxon.

We reconstructed significant cophylogenetic (coevolutionary) structure formed between weevils and their Nardonella endosymbionts. ParaFit test results rejected the null hypothesis that weevil and Nardonella phylogenies are similar due to chance alone (ParaFitGlobal = 138275, P = 0.00001) and supported significant global cophylogenetic structure between the hosts and their endosymbionts. Specifically, out of the 44 s between hosts and bacteria, 22 contributed to this global fit (Fig. tangletree). Early diverging clades of weevils, i.e., Brentidae, Dryophthorinae, and Entiminae, and their Nardonella endosymbionts show perfect cophylogenetic relationship. For the remaining clades, cophylogeny was observed mainly at “shallow” to “medium” levels (Fig. 4).

**Figure 3.**
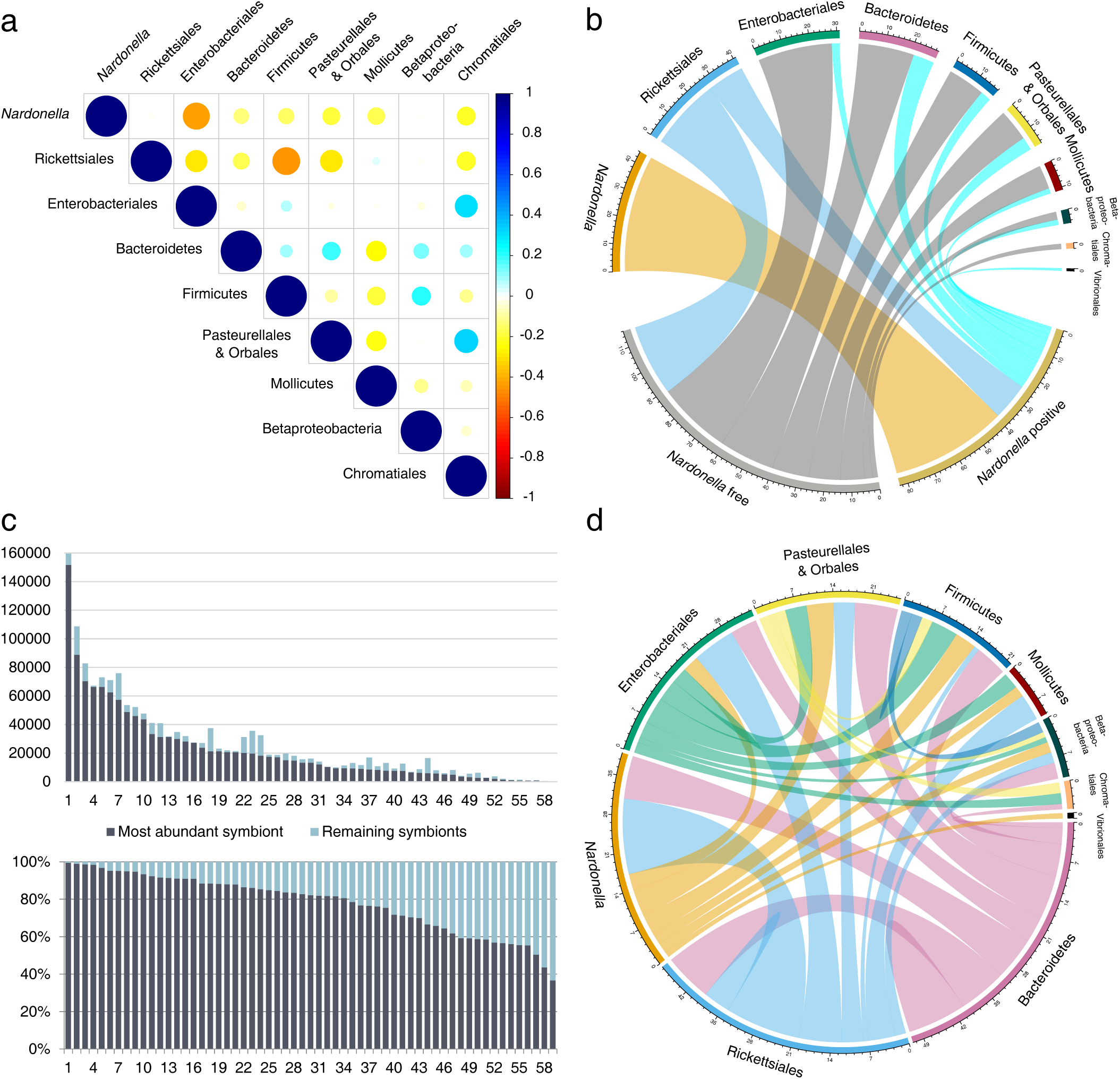
Correlation and coexistence among 10 major groups of endosymbionts.

**Figure 4.**
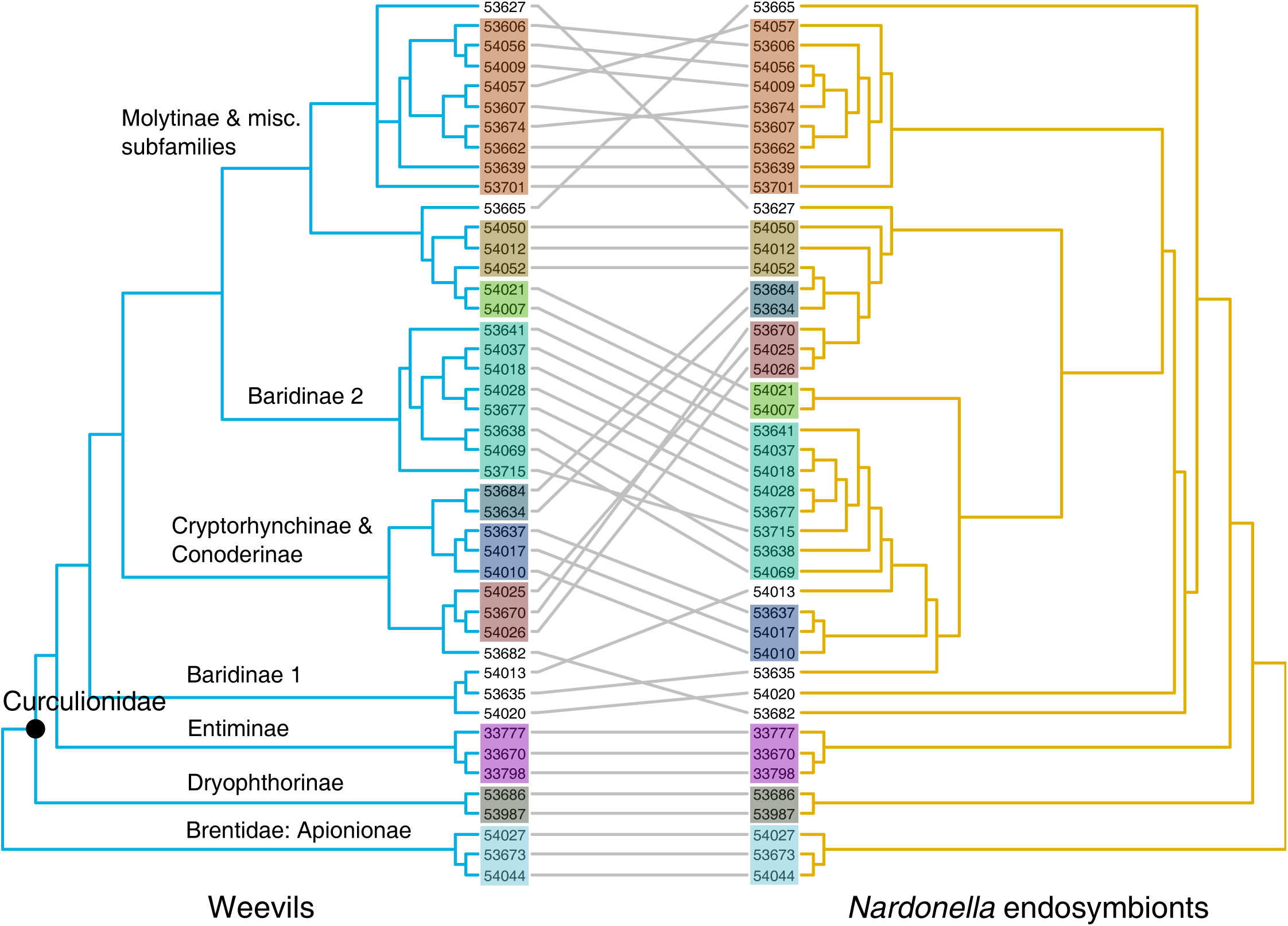
Cophylogeny between weevils and their *Nardonella* endosymbionts.

### Promiscuity, co-existence and co-exclusion

Fifty (45.9%) samples were infected with only one kind of endosymbiont, and 59 (54.1%) with 2 or more (38 with 2, 13 with 3, 7 with 4, and 1 with 6), thus considered as promiscuous samples. The majority or 75% (149/199) endosymbiont occurrences were associated with promiscuous samples. Endosymbionts belonging to Bacteroidetes exhibited co-existence 51 times, the highest among all, followed by Rickettsiales (48) and Nardonella (39) (Fig. Chord2). To adjust for differences in occurrence number between groups, the co-existence number was divided by the number of endosymbiont occurrences in a group, giving us a “co-existence ratio”. The lowest ratio (0.9 or 39/44) was found in Nardonella and Mollictues (0.91), and the highest in Betaproteobacteria (2.2) and Bacteriodetes (1.8) (excluding Chromatiales for low occurrence number). This means that on average each symbiont occurrence in the first two groups co-existed not more than once with another symbiont, whereas in the last two groups, it was about twice or more.

Overall, negative correlation between major endosymbiont groups was dominant, with 21 of the 36 possible pairs showing weak to moderate negative correlation (Spearman’s correlation coefficient between −0.1 to −0.5) (Fig. cormap). And those were concentrated in three groups, Nardonella, Rickettsiales and Mollicutes (Fig. cormap). Statistically significant, negative correlation was observed for the following pairs: Nardonella-Enterobacteriales (Spearman’s correlation coefficient=−0.42, P=0.0016), Rickettsiales-Firmicutes (-0.45, 0.0008), Rickettsiales-P & O (-0.29, 0.034), and Rickettsiales-Enterobacteriales (−0.28, 0.039). Statistically significant, positive correlation was observed only for Chromatiales-Enterobacteriales (0.3, 0.028) and Chromatiales-P & O (0.33, 0.016) (only two samples were infected with Chromatiales).

We will take a focused look at how endosymbiont occurrences sorted in relation to the presence of Nardonella. The distributions of Rickettsiales between Nardonella-free and Nardonella-positive samples were not statistically significant (chi-square test, χ^2^ = 0.85447, df = 1, P = 0.36). With Rickettsiales excluded, the distributions of the other 8 groups of endosymbionts were more strongly concentrated in Nardonella-free samples than in Nardonella-positive samples (chi-square test, df = 1, χ^2^ = 41.061, p-value ¡ 0.0001). The negative correlation or co-exclusion between Nardonella and Enterobacteriales is specially evident (chi-square test, df = 1, χ^2^ = 12.1, P ¡ 0.0005) (Fig. 3). Among the 31 occurrences of Enterobateriales (in 28 samples, as 3 samples each showed two occurrences), 28 were associated with Nardonella-free samples (n=69), and only 3 (9.7%) with Nardonella-positive samples (n=44) (Fig. Chord1). This means that samples hosting Nardonella were on average 6 times less likely to be infected with a bacterium in Enterobacteriales. This contrasts strongly with Rickettsiales—34.1% (14/41) of its occurrences co-existed with Nardonella-positive samples (Fig. 3).

